# All-in-One Adeno-associated Virus Delivery and Genome Editing by *Neisseria meningitidis* Cas9 *in vivo*

**DOI:** 10.1101/295055

**Authors:** Raed Ibraheim, Chun-Qing Song, Aamir Mir, Nadia Amrani, Wen Xue, Erik J. Sontheimer

## Abstract

Clustered, regularly interspaced, short palindromic repeats (CRISPR) and CRISPR-associated proteins (Cas) have recently opened a new avenue for gene therapy. Cas9 nuclease guided by a single-guide RNA (sgRNA) has been extensively used for genome editing. Currently, three Cas9 orthologs have been adapted for *in vivo* genome engineering applications: SpyCas9, SauCas9 and CjeCas9. However, additional *in vivo* editing platforms are needed, in part to enable a greater range of sequences to be accessed via viral vectors, especially those in which Cas9 and sgRNA are combined into a single vector genome. Here, we present an additional *in vivo* editing platform using *Neisseria meningitidis* Cas9 (NmeCas9). NmeCas9 is compact, edits with high accuracy, and possesses a distinct PAM, making it an excellent candidate for safe gene therapy applications. We find that NmeCas9 can be used to target the *Pcsk9* and *Hpd* genes in mice. Using tail vein hydrodynamic-based delivery of NmeCas9 plasmid to target the *Hpd* gene, we successfully reprogrammed the tyrosine degradation pathway in Hereditary Tyrosinemia Type I mice. More importantly, we delivered NmeCas9 with its single-guide RNA in a single recombinant adeno-associated vector (rAAV) to target *Pcsk9*, resulting in lower cholesterol levels in mice. This all-in-one vector yielded >35% gene modification after two weeks of vector administration, with minimal off-target cleavage *in vivo*. Our findings indicate that NmeCas9 can facilitate future efforts to correct disease-causing mutations by expanding the targeting scope of RNA-guided nucleases.

## Significance

We and others have shown that NmeCas9 is an efficient Cas9 nuclease with a high degree of specificity in cultured human cells. NmeCas9 has a unique PAM that expands the targeting range of Cas9-based genome editing. Since NmeCas9 is more compact than SpyCas9, its delivery with its guide-RNA in a single recombinant adeno-associated virus becomes a viable option. Furthermore, NmeCas9 has an unusually low off-target profile. These characteristics make NmeCas9 a strong candidate for *in vivo* genome editing, including therapeutic gene knockouts, as we demonstrate in the mouse liver. We anticipate that successful *in vivo* delivery of this accurate and effective Cas9 can advance therapeutic editing in humans.

## Introduction

A major advance in the field of gene therapy has been the introduction of Cas9 nuclease-enabled genome editing (1). Clustered, regularly interspaced, short palindromic repeats (CRISPR) loci specify an adaptive immune pathway that evolved in bacteria and archaea to defend against mobile genetic elements (MGEs) (2, 3). The effector complex in type II CRISPR systems includes the Cas9 nuclease, which is guided by a CRISPR RNA (crRNA) and a trans-activating RNA (tracrRNA). These dual RNAs can be fused to form a single-guide RNA (sgRNA) (4). Each crRNA contains a unique “spacer” sequence that can be programmed to cleave a DNA segment of interest. Cas9 scans DNA for a specific Protospacer Adjacent Motif (PAM), opens the duplex to form an RNA-DNA hybrid between the guide and the spacer, and introduces a double-strand break (DSB) in the DNA target (1, 3). Cas9 and sgRNA have been adapted to enable genome editing in cultured cells following various modes of delivery including plasmid and RNA transfections, viral transduction, and ribonucleoprotein (RNP) electroporation. Precise and efficient *in vivo* editing is more difficult to achieve, largely due to the difficulties inherent in delivery.

Several methods have been developed to deliver Cas9 *in vivo* including viral and non-viral methods (5). These include the use of gold and lipid nanoparticles to deliver Cas9 in RNP or RNA form in mice. However, these methods present challenges for routine use including cost and tissue distribution (6-8). One of the more intriguing gene delivery vehicles that has emerged in recent years is recombinant adeno-associated virus (rAAV). This vector possesses several attributes that benefit gene therapy applications, including lack of pathogenicity and replication as well as an ability to infect dividing and non-dividing cells (9). In addition, rAAV is also capable of infecting a wide range of cells and maintain sustained expression (10, 11). Compared to other viral vectors, rAAV persists in concatemeric, episomal forms, while eliciting mild immune responses (12-14). The usefulness of rAAV-based delivery for gene therapy is reflected in the number of clinical trials involving rAAV (15). One of the most exciting advancements for rAAV gene therapy field has been the FDA’s recent market approval of a therapy for RPE65-mediated inherited retinal disease (IRD), the first of its kind in the United States (16).

More recently, several groups have focused their efforts on using this tool for *in vivo* delivery of Cas9 orthologs (17-20). The majority of Cas9 genome editing efforts have been focused on the widely-used type II-A ortholog from *Streptococcus pyogenes*, SpyCas9. Although it exhibits consistently robust genome-editing activity, considerable effort has been required to overcome off-target editing activities of wild-type SpyCas9 (21-23) (Amrani *et al*., manuscript submitted). Furthermore, its large size (1,368 amino acids, 4.10 kb) restricts its delivery with its guide in a single virion with potent vectors such as rAAV (24). Split SpyCas9 constructs (expressed from separate viruses) have been employed (19), though activity is sometimes compromised (25-27). Dual-rAAV delivery of SpyCas9 and sgRNA can be achieved (28), but it requires the usage of highly minimized promoters that limit expression and tissue specificity. Furthermore, dual rAAV formats carry significant costs as well as limitations in co-transduction.

Alternatively, compact Cas9 orthologs can be packaged in all-in-one rAAV vectors. Type II-A *Staphylococcus aureus* (SauCas9) (1,053 amino acids, 3.16 kb) and type II-C *Campylobacter jejuni* Cas9 (CjeCas9) (984 amino acids, 2.95 kb) have been deployed successfully via rAAV in mice (18, 20). However, unlike the highly abundant NGG SpyCas9 PAM, these Cas9 nucleases have more restrictive PAM requirements (for SauCas9, 5’-NNGRRT-3’; for CjeCas9, 5’-N4RYAC (29). Furthermore, off-target editing by SauCas9 is not unusual (18, 30). For these reasons, many genomic sites of interest cannot be targeted by all-in-one rAAV delivery of the Cas9 genome editing machinery, and additional capabilities and PAM specificities are therefore needed.

We and others have reported genome editing in mammalian cells by the type II-C Cas9 from *Neisseria meningitidis* strain 8013 (NmeCas9) (31-33) [Amrani *et al*., manuscript submitted (http://www.biorxiv.org/content/early/2017/08/04/172650v2)]. NmeCas9 is small (1,082 amino acids, 3.16 kb), targets PAMs (N4GAYT > N4GYTT/N4GAYA/N4GTCT) that are distinct from those of the other compact Cas9 orthologs described above, and is intrinsically resistant to off-targeting (32) (Amrani *et al*., manuscript submitted). Additionally, NmeCas9 can be subjected to off-switch control by anti-CRISPR proteins (34), which could facilitate spatial and temporal control over NmeCas9 activity *in vivo* and *ex vivo*.

In this study, we report the *in vivo* delivery of NmeCas9 and its guide by a single expression cassette that is sufficiently small for all-in-one rAAV vectors. Two disease genes were targeted separately to highlight the therapeutic potential of NmeCas9: the *Hpd* gene in a hereditary tyrosinemia type I (HTI) mouse model (*Fah*^*neo*^), and the *Pcsk9* gene in C57Bl/6 mice. *Hpd* encodes the 4-hydroxyphenylpyruvate dioxygenase enzyme in the tyrosine metabolism pathway, and disrupting *Hpd* can lead to a decrease in the accumulation of toxic fumarylacetoacetate in tyrosinemia models (35). Separately, *Pcsk9* encodes proprotein convertase subtilisin/kexin type 9 (PCSK9), an antagonist of the low-density lipoprotein (LDL) receptor (36, 37). When PCSK9 is knocked out, more LDL receptors are available at the surface of hepatocytes to allow cholesterol binding and recycling towards the lysosomes for degradation (38, 39). The alleviation of tyrosinemia symptoms upon *Hpd* disruption, as well as the reduced serum cholesterol levels that result from *Pcsk9* disruption, provide convenient readouts for genome editing activity (18, 35). We used these systems to validate all-in-one rAAV delivery of NmeCas9 as an effective *in vivo* genome editing platform in mammals.

## Methods

### Construction of All-in-One AAV-sgRNA-hNMeCas9 Plasmid and rAAV vector production

The human-codon-optimized NmeCas9 gene under the control of the U1a promoter, and a single-guide RNA cassette driven by the U6 promoter, were cloned into an AAV2 plasmid backbone. The NmeCas9 ORF was flanked by four nuclear localization signals – two on each terminus – in addition to a triple-HA epitope tag. Oligonucleotides with spacer sequences targeting *Hpd, Pcsk9, and Rosa26* were inserted into the sgRNA cassette by ligation into a SapI cloning site (**Supplementary Note**).

AAV vector production was performed at the Horae Gene Therapy Center at the University of Massachusetts Medical School. Briefly, plasmids were packaged in AAV8 capsid by triple-plasmid transfection in HEK 293 cells and purified by sedimentation as previously described (40).

The off-target profiles of these spacers were predicted computationally using the Bioconductor package CRISPRseek. Search parameters were adapted to NmeCas9 settings as described previous (Amrani *et al*., manuscript submitted): gRNA.size = 24, PAM = “NNNNGATT”, PAM.size = 8, RNA.PAM.pattern = “NNNNGNNN$”, weights = c(0, 0, 0, 0, 0, 0, 0.014, 0, 0, 0.395, 0.317, 0, 0.389, 0.079, 0.445, 0.508, 0.613, 0.851, 0.732, 0.828, 0.615, 0.804, 0.685, 0.583), max.mismatch = 6, allowed.mismatch.PAM = 7, topN = 10000, min.score = 0.

### Cell culture and transfection

Mouse Hepa 1-6 hepatoma cells were cultured in DMEM with 10% FBS and 1% Penicillin/Streptomycin (Gibco) in a 37 °C incubator with 5% CO2. Transient transfections of Hepa 1-6 cells were performed using Lipofectamine LTX. For transient transfection, approximately 1×10^5^ cells per well were cultured in 24-well plates 24 hours before transfection. Each well was transfected with 500 ng all-in-one AAV-sgRNA-hNmeCas9 plasmid, using Lipofectamine LTX with Plus Reagent (Invitrogen) according to the manufacturer’s protocol.

### DNA isolation from cells and liver tissue

Genomic DNA isolation from cells was performed 72 hours post-transfection cells using DNeasy Blood and Tissue kit (Qiagen) following the manufacturer’s protocol. Mice were sacrificed and liver tissues were collected 10 days post-hydrodynamic injection or 14 and 50 days post-tail vein rAAV injection. Genomic DNA was isolated using DNeasy Blood and Tissue kit (Qiagen) according to the manufacturer’s protocol.

### GUIDE-seq

GUIDE-seq analysis was performed as previously described (22). Briefly, 7.5 pmol of annealed GUIDE-seq oligonucleotides and 500 ng of all-in-one AAV-sgRNA-hNmeCas9 plasmids targeting *Pcsk9, Rosa26* and *Hpd* were transfected into 1×10^5^ Hepa 1-6 cells using Lipofectamine LTX with Plus Reagent (Invitrogen). At 72 hours post-transfection, genomic DNA was extracted using a DNeasy Blood and Tissue kit (Qiagen) per manufacturer protocol. Library preparations, deep sequencing, and reads analysis were performed as previously described (41, 42). The Bioconductor package GUIDEseq was used for off-target analysis as described previously using maximum allowed mismatch of 10 nt between the guide and target DNA (41). For read alignment, mouse mm10 was used as a reference genome.

### Indel analysis

TIDE primers were designed ~700 bp apart, with the forward primer at ~200 bp upstream of the cleavage site. 50 ng of genomic DNA was used for PCR amplification with High Fidelity 2X PCR Master Mix (New England Biolabs). For TIDE analysis, 30 μl of a PCR product was purified using QIAquick PCR Purification Kit (Qiagen) and sent for Sanger sequencing using the TIDE forward primer (**Supplementary Table**). Indel values were obtained using the TIDE web tool (https://tide-calculator.nki.nl/) as described previously (43).

Targeted deep-sequencing analysis was performed for Hepa 1-6 cells and mouse liver gDNA using a two-step PCR amplification approach as described previously (42) (Amrani *et al*., manuscript submitted). Briefly, in the first PCR step, on-target or off-target locus-specific primers were used to amplify the editing site using Phusion High Fidelity DNA Polymerase (New England Biolabs) with a 65°C annealing temperature. The primer ends contained sequences complementary to Illumina TruSeq adaptor sequences (**Supplementary Table**). In the second-step PCR, equimolar amounts of DNA were amplified with a universal forward primer and an indexed reverse primer using Phusion High Fidelity DNA Polymerase (98 °C, 15s; 61°C, 25s; 72 °C, 18s; 9 cycles) to ligate the TruSeq adaptors. The resultant amplicons were separated in a 2.5% agarose gel and the corresponding ~250 bp product bands were extracted using Monarch DNA Gel Extraction Kit (New England Biolabs).

The libraries were then sequenced on an Illumina MiSeq in paired-end mode with a read length of 150 bp. To analyze genome editing outcomes at genomic sites, the command line utilities of CRISPResso were used (44). Input parameters were adjusted to filter low-quality reads (-q 30 -s 20). Furthermore, the background was determined using the control sample (no guide) and subtracted from the experimental samples. The resulting indel frequencies, sizes and distributions were then plotted using Graphpad PRISM.

### Animals and liver tissue processing

All animal procedures were reviewed and approved by The Institutional Animal Care and Use Committee (IACUC) at University of Massachusetts Medical School. For hydrodynamic injections, 2.5 mL of 30 μg of endotoxin-free AAV-sgRNA-hNmeCas9 plasmid targeting *Pcsk9* or 2.5 ml PBS was injected by tail-vein into 9- to 18-week-old female C57BL/6 mice. Mice were euthanized 10 days later, and liver tissue was harvested. For the AAV8 vector injections, 12- to 16-week-old female C57BL/6 mice were injected with 4 x10^11^ genome copies per mouse via tail vein, using vectors targeting *Pcsk9* or *Rosa26*. Mice were sacrificed 14 and 50 days after vector administration and liver tissues were collected for analysis.

For *Hpd* targeting, 2 mL PBS or 2 mL of 30 μg of endotoxin-free AAV-sgRNA-hNmeCas9 plasmid was administered into 15-to 21-week-old Type 1 Tyrosinemia *Fah* knockout mice (*Fah^neo^*) via tail vein. The encoded sgRNAs targeted sites in exon 8 (*sgHpd1*) or exon 11 (*sgHpd2*). The HT1 homozygous mice with the *Fah^neo^* allele in a 129 background were kindly provided by Dr. Markus Grompe (45). The HT1 mice were fed with 10 mg/L NTBC [2-(2-nitro-4-trifluoromethylbenzoyl)-1,3-cyclohexanedione] (Sigma-Aldrich, Cat. No. PHR1731-1G) in drinking water when indicated. Both sexes were used in these experiments. Mice were maintained on NTBC water for 7 days postinjection and then switched to normal water. Body weight was monitored every 1-3 days. The PBS-injected control mice were sacrificed when they became moribund after losing 20% of their body weight after removal from NTBC treatment.

Mice were euthanized according to our protocol and liver tissue was sliced and fragments stored at −80°C. Some liver tissues were fixed in 4% formalin overnight, embedded in paraffin, sectioned and stained with hematoxylin and eosin (H&E).

### Serum analysis

Blood (~200 μL) was drawn from the facial vein at 0, 25 and 50 days post vector administration. Serum was isolated using a serum separator (BD, Cat. No. 365967) and stored under −80 °C until assay.

Serum cholesterol levels were measured by Infinity™ colorimetric endpoint assay (Thermo-Scientific) following the manufacturer’s protocol. Briefly, serial dilutions of Data-Cal™ Chemistry Calibrator were prepared in PBS. In a 96-well plate, 2 μL of mice sera or calibrator dilution was mixed with 200 μL of Infinity™ cholesterol liquid reagent, then incubated at 37 °C for 5 minutes. The absorbance was measured at 500 nm using a BioTek Synergy HT microplate reader.

### Western blot

Liver tissue fractions were ground and resuspended in 150 μL of RIPA lysis buffer. Total protein content was estimated by Pierce™ BCA Protein Assay Kit (Thermo-Scientific) following the manufacturer’s protocol. 20 μg of protein from tissue or 2 ng of Recombinant Mouse Proprotein Convertase 9/PCSK9 Protein (R&D Systems, 9258-SE-020) were loaded onto a 4-20% Mini-PROTEAN^®^ TGX™ Precast Gel (Bio-Rad). The separated bands were transferred onto PVDF membrane and blocked with 5% Blocking-Grade Blocker solution (Bio-Rad) for 2 hours at room temperature. Membranes were incubated with rabbit anti-GAPDH (Abcam ab9485, 1:2000) or goat anti-PCSK9 (R&D Systems AF3985, 1:400) antibodies overnight at 4 °C. Membranes were washed five times in TBST and incubated with horseradish peroxidase (HRP)-conjugated goat anti-rabbit (Bio-Rad 1706515, 1:4000), and donkey anti-goat (R&D Systems HAF109, 1:2000) secondary antibodies for 2 hours at room temperature. The membranes were washed five times in TBST and visualized with Clarity™ western ECL substrate (Bio-Rad) using an M35A X-OMAT Processor (Kodak).

### Humoral Immune Response

Humoral IgG1 immune response to NmeCas9 was measured by ELISA (Bethyl; Mouse IgG1 ELISA Kit, E99-105) following manufacturer’s protocol with a few modifications. Briefly, expression and three-step purification of NmeCas9 and SpyCas9 was performed as previously described (4). 0.5 μg of recombinant NmeCas9 or SpyCas9 proteins suspended in 1x coating buffer (Bethyl) were used to coat 96-well plates (Corning), and incubated for 12 hours at 4 °C with shaking. The wells were washed 3 times while shaking for 5 minutes using 1x Wash Buffer. Plates were blocked with 1x BSA Blocking Solution (Bethyl) for 2 hours at room temperature, then washed three times. Serum samples were diluted 1:40 using PBS and added to each well in duplicate. After incubating the samples at 4 °C for 5 hours, the plates were washed 3x times for 5 minutes and 100 μL of biotinylated anti-mouse IgG1 antibody (Bethyl; 1: 100,000 in 1 x BSA Blocking Solution) was added to each well. After incubating for 1 hour at room temperature, the plates were washed 4 times, and 100 μL of TMB Substrate was added to each well. The plates were allowed to develop in the dark for 20 minutes at room temperate, and 100 μL of ELISA Stop Solution was then added per well. Following the development of the yellow solution, absorbance was recorded at 450 nm using a BioTek Synergy HT microplate reader.

## Results

### Efficient Genome Editing Using All-in-One AAV-sgRNA-hNmeCas9 Plasmid in Cells and *in vivo* by Hydrodynamic Injection

Recently, we have shown that the relatively compact NmeCas9 is active in genome editing in a range of cell types (Amrani *et al*., manuscript submitted). To exploit the small size of this Cas9 ortholog, we generated an all-in-one AAV construct with human-codon-optimized NmeCas9 under the expression of the mouse U1a promoter, and with its sgRNA driven by the U6 promoter (**Figure 1A; Supplementary Note**).

**Figure 1:**
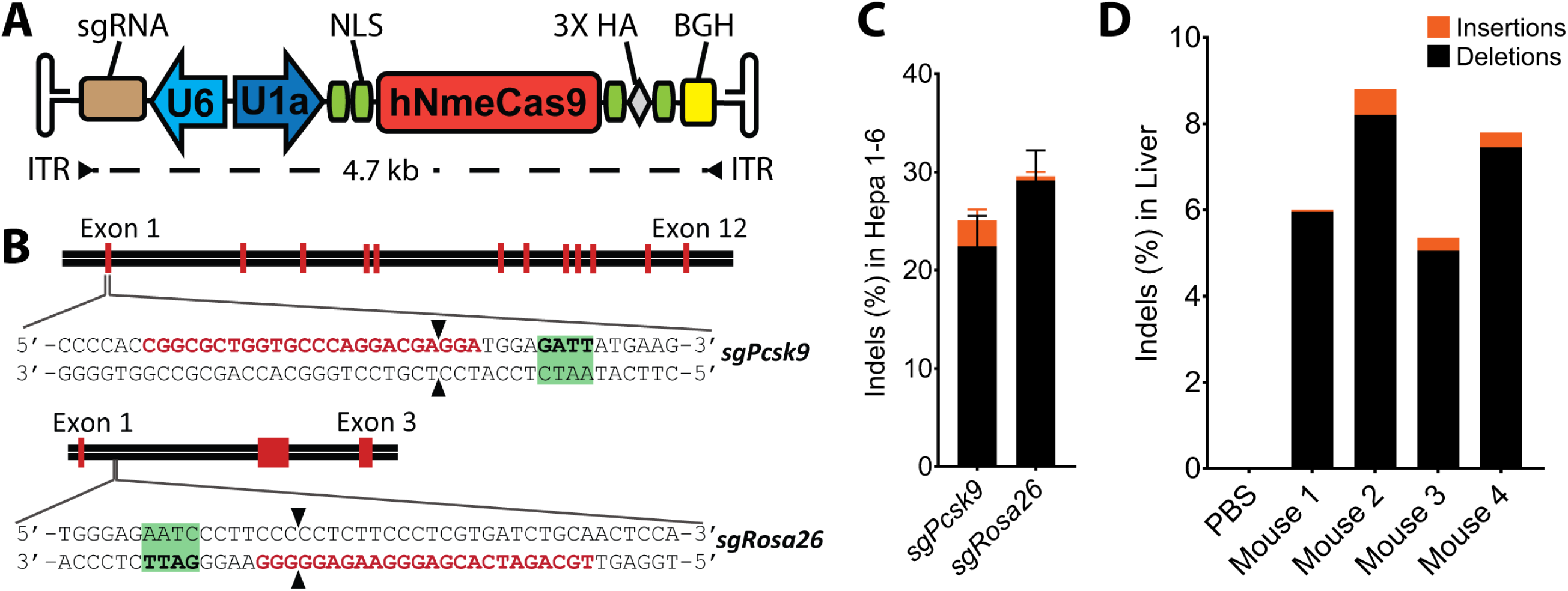
Validation of an all-in-one AAV-sgRNA-hNmeCas9 construct. **(A)** Schematic representation of a single rAAV vector expressing human-codon-optimized NmeCas9 and its sgRNA. The backbone is flanked by AAV inverted terminal repeats (ITR). The poly(A) signal is from rabbit beta-globin (BGH). **(B)** Schematic diagram of the *Pcsk9* (top) and *Rosa26* (bottom) mouse genes. Red bars represent exons. Zoomed-in views show the protospacer sequence (red) whereas the NmeCas9 PAM sequence is highlighted in green. Double-stranded break location sites are denoted (black arrowheads). **(C)** Stacked histogram showing the percentage distribution of insertions-deletions (indels) obtained by TIDE after AAV-sgRNA-hNmeCas9 plasmid transfections in Hepa 1–6 cells targeting *Pcsk9* (*sgPcsk9*) and *Rosa26* (*sgRosa26*) genes. Data are presented as mean values ± SD from three biological replicates. **(D)** Stacked histogram showing the percentage distribution of indels at *Pcsk9* in the liver of C57Bl/6 mice obtained by TIDE after hydrodynamic injection of AAV-sgRNA-hNmeCas9 plasmid.

Two sites in the mouse genome were selected initially to test the nuclease activity of NmeCas9 *in vivo*: the *Rosa26* “safe-harbor” gene (targeted by *sgRosa26*), and the proprotein convertase subtilisin/kexin type 9 (*Pcsk9*) gene (targeted by *sgPcsk9*), a common therapeutic target for lowering circulating cholesterol and reducing the risk of cardiovascular disease (**Figure 1B**). Genome-wide off-target predictions for these guides were determined computationally using the Bioconductor package CRISPRseek 1.9.1 (46) with N_4_GN_3_ PAMs and up to 6 mismatches. Many N_4_GN_3_ PAMS are inactive, so these search parameters are nearly certain to cast a wider net than the true off-target profile. Despite the expansive nature of the search, our analyses revealed no off-target sites with fewer than four mismatches in the mouse genome (**Supplementary Figure 1**). On-target editing efficiencies at these target sites were evaluated in mouse Hepa 1-6 hepatoma cells by plasmid transfections and indel quantification was performed by sequence trace decomposition using the TIDE web tool. We found >25% indel values for the selected guides, the majority of which were deletions (**Figure 1C**).

To evaluate the preliminary efficacy of the constructed all-in-one AAV-sgRNA-hNmeCas9 vector, endotoxin-free *sgPcsk9* plasmid was hydrodynamically administered into the C57Bl/6 mice via tail-vein injection. This method can deliver plasmid DNA to ~40% of hepatocytes for transient expression (47). Indel analyses by TIDE using DNA extracted from liver tissues revealed 5-9% indels 10 days after vector administration (**Figure 1D**), comparable to the editing efficiencies obtained with analogous tests of SpyCas9 (48). These results suggested that NmeCas9 is capable of editing liver cells *in vivo*.

### Knockout of 4-Hydroxyphenylpyruvate Dioxygenase Rescues the Lethal Phenotypes of Hereditary Tyrosinemia Type I Mice

Hereditary Tyrosinemia type I (HT-I) is a fatal genetic disease caused by autosomal recessive mutations in the *Fah* gene, which codes for the fumarylacetoacetate hydroxylase (FAH) enzyme. Patients with diminished FAH have a disrupted tyrosine catabolic pathway, leading to the accumulation of toxic fumarylacetoacetate and succinyl acetoacetate, causing liver and kidney damage (49). Over the past two decades, the disease has been controlled by 2-(2-Nitro-4-trifluoromethylbenzoyl) -1,3-cyclohexanedione (NTBC), which inhibits 4-hydroxyphenylpyruvate dioxygenase upstream in the tyrosine degradation pathway, thus preventing the accumulation of the toxic metabolites (50). However, this treatment requires lifelong management of diet and medication, and may eventually require liver transplantation (51).

Several gene therapy strategies have been tested to correct the defective *Fah* gene using site-directed mutagenesis (52) or homology-directed repair by CRISPR-Cas9 (52-54). It has been reported that successful modification of only 1/10,000 of hepatocytes in the liver is sufficient to rescue the phenotypes of *Fah^mut/mut^* mice. Recently, a metabolic pathway reprogramming approach has been suggested in which the function of the hydroxyphenylpyruvate dioxygenase (HPD) enzyme was disrupted by the deletion of exons 3 and 4 of the *Hpd* gene in the liver (35). This provides us with a context in which to test the efficacy of NmeCas9 editing, by targeting *Hpd* and assessing rescue of the disease phenotype in *Fah* mutant mice (45). For this purpose, we screened and identified two target sites [one each in exon 8 (*sgHpd1*) and exon 11 (*sgHpd2*)] within the open reading frame of *Hpd* (**Figure 2A**). These guides induced average indel efficiencies of 10.8% and 9.1%, respectively by plasmid transfections in Hepa 1-6 cells (**Supplementary Figure 2**).

**Figure 2:**
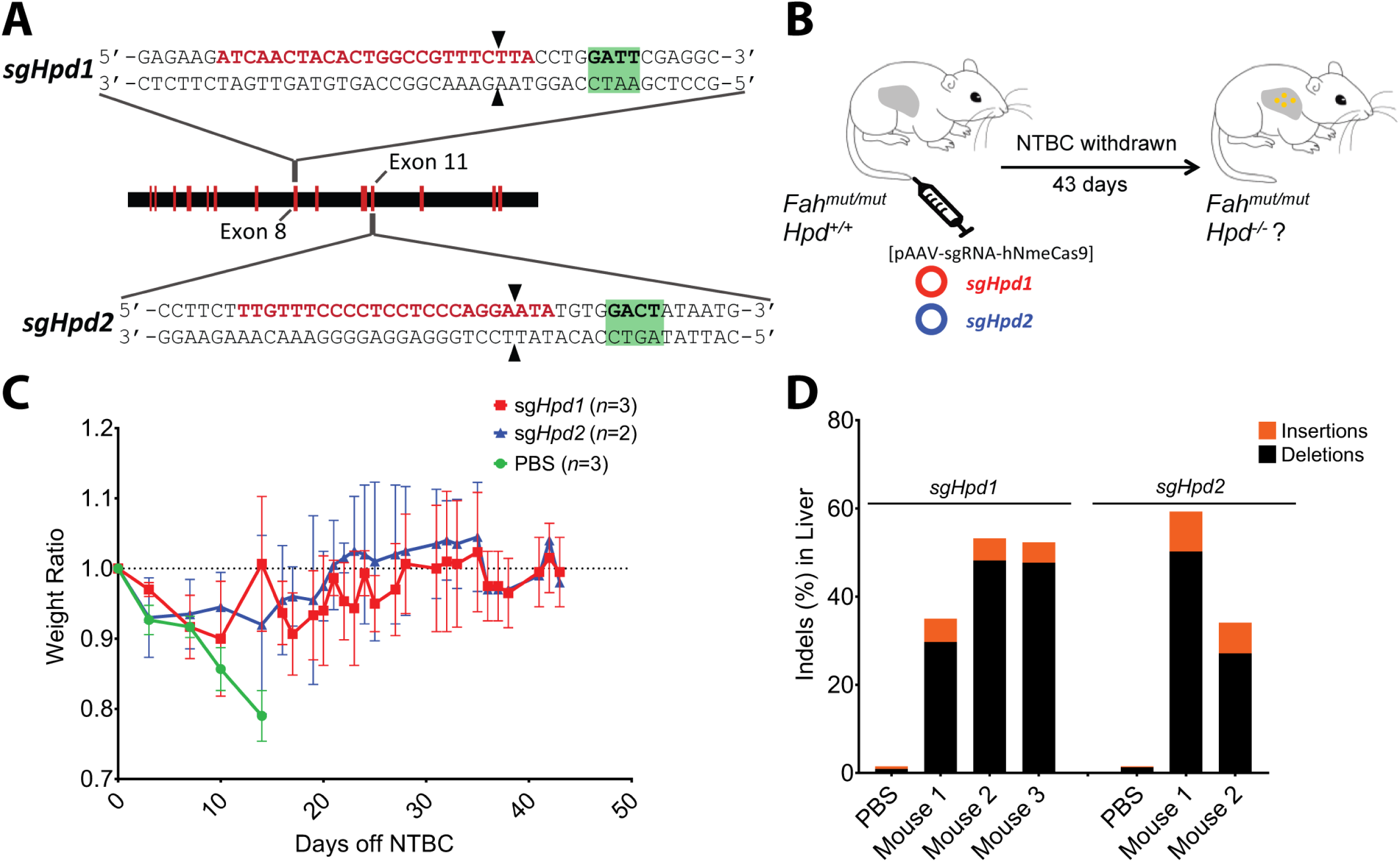
NmeCas9-mediated knockout of *Hpd* rescues the lethal phenotype in Hereditary Tyrosinemia Type I Mice. **(A)** Schematic diagram of the *Hpd* mouse gene. Red bars represent exons. Zoomed-in views show the protospacer sequences (red) for targeting exon 8 (*sgHpd1*) and exon 11 (*sgHpd2*). NmeCas9 PAM sequences are in green, and double-stranded break locations are indicated (black arrowheads). **(B)** Experimental design. Three groups of Hereditary Tyrosinemia Type I *Fah^-/-^* mice are injected with PBS, or all-in-one AAV-sgRNA-hNmeCas9 plasmids *sgHpd1* or *sgHpd2*. **(C)** Weight of mice hydrodynamically injected with PBS (green), AAV-sgRNA-hNmeCas9 plasmid *sgHpd1* targeting *Hpd* exon 8 (red) or *sgHpd2* targeting *Hpd* exon 11 (blue) were monitored after NTBC withdrawal. Error bars represent three mice for PBS and *sgHpd1* groups, and two mice for the *sgHpd2* group. Data are presented as mean ± SD. **(D)** Stacked histogram showing the percentage distribution of indels at *Hpd* in liver of *Fah^-/-^* mice obtained by TIDE after hydrodynamic injection of PBS, or *sgHpd1* and *sgHpd2* plasmids. Livers were harvested at the end of NTBC withdrawal (day 43).

Three groups of mice were treated by hydrodynamic injection with either PBS, or with one of the two *sgHpd1* and *sgHpd2* all-in-one AAV-sgRNA-hNmeCas9 plasmids. One mouse in the *sgHpd1* group and two in the *sgHpd2* group were excluded from the follow-up study due to failed tail-vein injections. Mice were taken off NTBC-containing water 7 days after injections, and their weight was monitored for 43 days post-injection (**Figure 2B**). Mice injected with PBS suffered severe weight loss (a hallmark of HT-I), and were sacrificed after losing 20% of their body weight (**Figure 2C**). Overall, all *sgHpd1* and *sgHpd2* mice successfully maintained their body weight for 43 days overall, and for at least 21 days without NTBC (**Figure 2C**). NTBC treatment had to be resumed for 2-3 days for two mice that received *sgHpd1* and one that received *sgHpd2* to allow them to regain body weight during the 3^rd^ week after plasmid injection, perhaps due to low initial editing efficiencies, liver injury due to hydrodynamic injection, or both. Conversely, all other *sgHpd1* and *sgHpd2* treated mice achieved indels with frequencies ranging from 35% to 60% (**Figure 2D**). This level of gene inactivation likely reflects not only the initial editing events but also the competitive expansion of edited cell lineages (after NTBC withdrawal) at the expense of their unedited counterparts (51, 52, 54). Liver histology revealed that liver damage is substantially less severe in the *sgHpd1* and *sgHpd2* treated mice compared to *Fah^mut/mut^* mice injected with PBS, as indicated by the smaller numbers of multinucleated hepatocytes compared to PBS-injected mice (**Supplementary Figure 3**).

### *In vivo* Genome Editing by NmeCas9 Delivered by a rAAV Vector

Although plasmid hydrodynamic injections can generate indels, therapeutic development will require less invasive delivery strategies, such as rAAV. To this end, all-in-one AAV-sgRNA-hNmeCas9 plasmids were packaged in hepatocyte-tropic AAV8 capsids to target *Pcsk9* (*sgPcsk9*) and *Rosa26* (*sgRosa26*) (**Figure 1B**) (55, 56). Vectors were administered into C57BL/6 mice via tail vein (**Figure 3A**). We monitored cholesterol level in the serum, and measured PCSK9 protein and indel frequencies in the liver tissues 25 and 50 days post injection.

**Figure 3:**
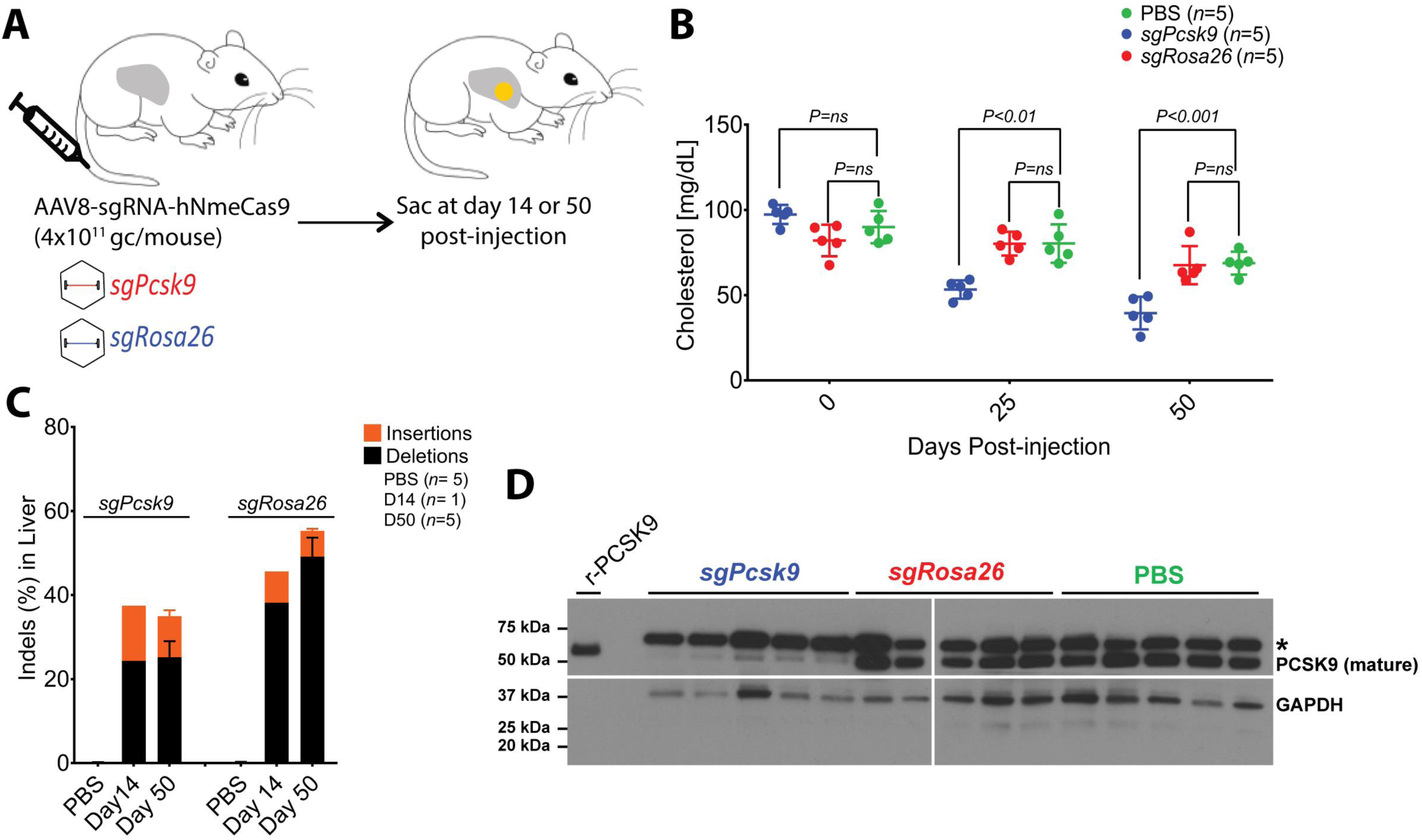
AAV-delivery of NmeCas9 for *in vivo* genome editing. **(A)** Experimental outline of AAV8-sgRNA-hNmeCas9 vector tail-vein injections to target *Pcsk9* (*sgPcsk9*) and *Rosa26* (*sgRosa26*) in C57Bl/6 mice. Mice were sacrificed at 14 (n = 1) or 50 days (n = *5*) post-injection and liver tissues were harvested. Blood sera were collected at day 0, 25 and 50 post-injection for cholesterol level measurement. **(B)** Serum cholesterol levels. *P* values are calculated by unpaired t test. **(C)** Stacked histogram showing the percentage distribution of indels at *Pcsk9* or *Rosa26* in livers of mice, as measured by targeted deep-sequencing analyses. Data are presented as mean ± SD from five mice per cohort. **(D)** A representative anti-PCSK9 western blot using total protein collected from day 50 mouse liver homogenates. 2 ng of recombinant mouse PCSK9 (r-PCSK9) was included as a mobility standard. The asterisk indicates a cross-reacting protein that is larger than the control recombinant protein.

Using a colorimetric endpoint assay, we determined that the circulating serum cholesterol level in the *sgPcsk9* mice decreased significantly (*p* < 0.001) compared to the PBS and *sgRosa26* mice at 25 and 50 days post-injection (**Figure 3B**). Targeted deep-sequencing analyses at *Pcsk9* and *Rosa26* target sites revealed very efficient indels of 35% and 55% respectively at 50 days post-vector administration (Figure 3C). Additionally, one mouse of each group was euthanized at 14 days post-injection, and revealed on-target indel efficiencies of 37% and 46% at *Pcsk9* and *Rosa26*, respectively (**Figure 3C**). As expected, PCSK9 protein levels in the livers of *sgPcsk9* mice were substantially reduced compared to the mice injected with PBS and *sgRosa26* (**Figure 3D**). The efficient editing, PCSK9 reduction, and diminished serum cholesterol indicate the successful delivery and activity of NmeCas9 at the *Pcsk9* locus.

SpyCas9 delivered by viral vectors is known to elicit host immune responses (19, 57). To investigate if the mice injected with AAV8-sgRNA-hNmeCas9 generate anti-NmeCas9 antibodies, we used sera from the treated animals to perform IgG1 ELISA. Our results show that NmeCas9 elicits a humoral response in these animals (**Supplementary Figure 4**). Despite the presence of an immune response, NmeCas9 delivered by rAAV is highly functional *in vivo*, with no apparent signs of abnormalities or liver damage (**Supplementary Figure 5**).

**Figure 4:**
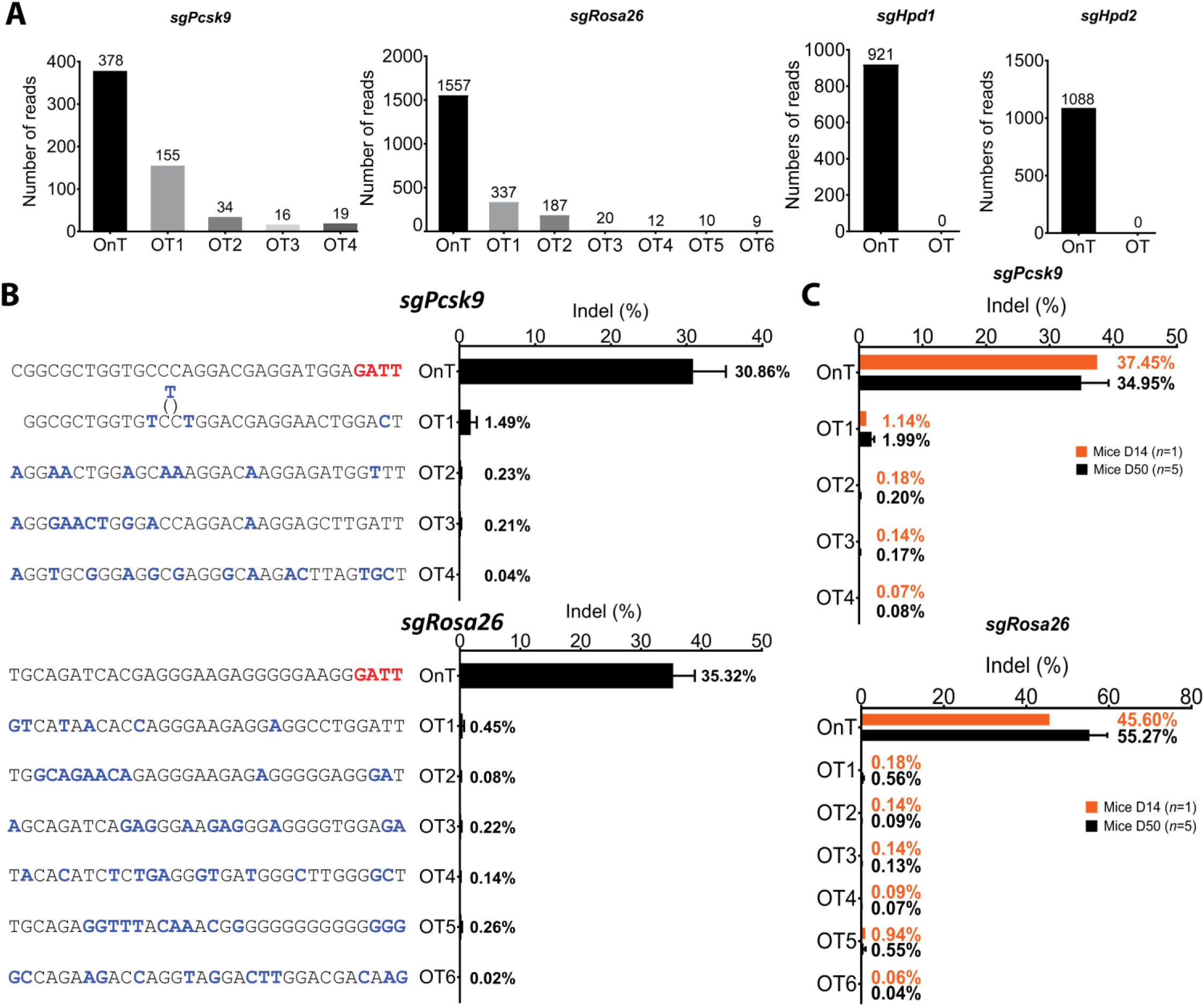
GUIDE-seq genome-wide specificities of NmeCas9. **(A)** Number of GUIDE-seq reads for the on-target (OnT) and off-target (OT) sites. **(B)** Targeted deep-sequencing to measure the lesion rates at each of the OT sites in Hepa 1–6 cells. The mismatches of each OT site with the OnT protospacers is highlighted (blue). Data are presented as mean ± SD from three biological replicates. **(C)** Targeted deep sequencing to measure the lesion rates at each of the OT sites using genomic DNA obtained from mice injected with all-in-one AAV8-sgRNA-hNmeCas9 *sgPcsk9* and *sgRosa26* and sacrificed at day 14 (D14) or day 50 (D50) post-injection. Data are presented as mean ± SD.

### NmeCas9 is Highly Specific *in vivo*

A significant concern in therapeutic CRISPR/Cas9 genome editing is the possibility of off-target edits. We and others have found that wildtype NmeCas9 is a naturally high-accuracy genome editing platform in cultured mammalian cells (32) (Amrani *et al*., manuscript submitted). To determine if NmeCas9 maintains its minimal off-targeting profile in mouse cells and *in vivo*, we screened for off-target sites in the mouse genome using genome-wide, unbiased identification of DSBs enabled by sequencing (GUIDE-seq) (22). Hepa 1-6 cells were transfected with *sgPcsk9, sgRosa26, sgHpd1* and *sgHpd2* all-in-one AAV-sgRNA-hNmeCas9 plasmids and the resulting genomic DNA was subjected to GUIDE-seq analysis. Consistent with our previous observations in human cells (Amrani *et al*., manuscript submitted), GUIDE-seq revealed very few off-target (OT) sites in the mouse genome. Four potential OT sites were identified for *sgPcsk9* and another six for *sgRosa26*. We were unable to detect off-target edits with *sgHpd1* and *sgHpd2* (**Figure 4A**), thus reinforcing our previous observation that NmeCas9 is often intrinsically hyper-accurate (Amrani *et al*., manuscript submitted).

Several of the putative OT sites for *sgPcsk9* and *sgRosa26* lack the NmeCas9 PAM preferences (N_4_GATT, N_4_GCTT, N_4_GTTT, N_4_GACT, N_4_GATA, N_4_GTCT, and N_4_GACA) (**Figure 4B**) and may therefore represent background. To validate these OT sites, we performed targeted deep-sequencing using genomic DNA from Hepa 1-6 cells. By this more sensitive readout, indels were undetectable above background at all these OT sites except OT1 of *Pcsk9*, which had an indel frequency less than 2% (Figure 4B). To validate NmeCas9’s high fidelity *in vivo*, we measured indel formation at these OT sites in liver genomic DNA from the AAV8-NmeCas9-treated, *sgPcsk9-* and *sgRosa26-*targeted mice. We found little or no detectable off-target editing in mice liver sacrificed at 14 days at all sites except *sgPcsk9* OT1, which exhibited fewer than 2% lesion efficiency (**Figure 4C**). More importantly, this level of OT editing stayed below <2% even after 50 days, and also remained either undetectable or very low for all other candidate OT sites. These results suggested that extended (50 days) expression of NmeCas9 *in vivo* does not compromise its targeting fidelity (**Figure 4C**).

## Discussion

### All-in-One rAAV delivery of hNmeCas9

Compared to transcription activator-like effector nucleases (TALENs) and Zinc-finger nucleases (ZFNs), Cas9s are distinguished by their flexibility and versatility (1). Such characteristics make them ideal for driving the field of genome engineering forward. Over the past few years, CRISPR-Cas9 has been used to enhance products in agriculture, food and industry, in addition to the promising applications in gene therapy and personalized medicine (58). Despite the diversity of Class 2 CRISPR systems that have been described, only a handful of them have been developed and validated for genome editing *in vivo*. In this study, we have shown that NmeCas9 is a compact, high-fidelity Cas9 that can be considered for future *in vivo* genome editing applications using all-in-one rAAV. Its unique PAM enables editing at additional targets that are inaccessible to the other two compact, all-in-one rAAV-validated orthologs (SauCas9 and CjeCas9).

### Therapeutic Gene Correction for Hereditary Tyrosinemia Type 1 by Metabolic Pathway Reprograming

Patients with mutations in the *HPD* gene are considered to have Type III Tyrosinemia and exhibit high level of tyrosine in blood, but otherwise appear to be largely asymptomatic (59, 60). HPD acts upstream of FAH in the tyrosine catabolism pathway, and *Hpd* disruption ameliorates HT-I symptoms by preventing the toxic metabolite buildup that results from loss of FAH. Structural analyses of HPD reveal that the catalytic domain of the HPD enzyme is located at the C-terminus of the enzyme and encoded by exon 13 and 14 (61). Thus, frameshift-inducing indels upstream of exon 13 should render the enzyme inactive. We used this context to demonstrate that *Hpd* inactivation by hydrodynamic injection of NmeCas9 plasmid is a viable approach to rescue HT-I mice. NmeCas9 can edit sites carrying several different PAMs [N_4_GATT (consensus), N_4_GCTT, N_4_GTTT, N_4_GACT, N_4_GATA, N_4_GTCT, and N_4_GACA] (Amrani *et al*., manuscript submitted). Our *Hpd* editing experiments confirmed one of the variant PAMs *in vivo* with the *sgHpd2* guide, which targets a site with a N_4_GACT PAM.

### Efficient, Accurate NmeCas9 Genome Editing with rAAV Delivery

To achieve targeted delivery of NmeCas9 to various tissues *in vivo*, rAAV vectors are a promising delivery platform due to the compact size of NmeCas9 transgene, which allows the delivery of NmeCas9 and its guide in all-in-one format. We have validated this approach for the targeting of *Pcsk9* and *Rosa26* genes in adult mice, with efficient editing observed even at 14 days postinjection. As observed previously in cultured cells (32) (Amrani *et al*., manuscript submitted), NmeCas9 is intrinsically accurate, even without the extensive engineering that was required to reduce off-targeting by SpyCas9 (21-23). We performed side-by-side comparisons of NmeCas9 OT editing in cultured cells and *in vivo* by targeted deep-sequencing, and we found that off-targeting is minimal in both settings. Editing at the *sgPcsk9* OT1 site (within an unannotated locus) was the highest detectable at ~2%. Despite these promising results, more extensive and long-term studies, including in larger animals, will be needed to fully understand the long-term effects of Cas9 expression in tissues, as well as the development of approaches that clear viral vectors after editing is complete.

In conclusion, we demonstrate that NmeCas9 is amenable to *in vivo* genome editing using the highly desirable all-in-one rAAV platform. With its unique PAM preferences and high fidelity, this all-in-one AAV-sgRNA-hNeCas9 can be applied to a range of genome editing purposes *in vivo*.

## Acknowledgments

We thank Haiwei Mou and Yueying Cao for assistance with mouse injections, and Xin Gao for assistance in designing of deep sequencing and GUIDE-seq libraries. We also thank the UMMS Viral Vector Core for AAV packaging services, and the UMMS Deep Sequencing Core for sequencing. We are grateful to Guangping Gao, Scot Wolfe, and all members of the Xue and Sontheimer labs for discussions, advice, and helpful feedback. W.X. was supported by the NIH (DP2HL137167 and P01HL131471), the Lung Cancer Research Foundation, the ALS Association, the American Cancer Society (129056-RSG-16-093), and Hyundai Hope on Wheels. E.J.S. was supported by the NIH (R01GM1115911 to Scot A. Wolfe and E.J.S.). E.J.S. is a co-founder and advisor of Intellia Therapeutics, Inc.

## Author Contributions

R.I. and E.J.S. initiated the study, and R.I., C.S., W.X. and E.J.S. designed experiments. R.I. designed, constructed, and validated all-in-one AAV-sgRNA-hNmeCas9 plasmids in cells, prepared GUIDE-seq and targeted lesion libraries from cultured cells and mouse tissues, and analyzed *in vivo*-edited samples. C.S. performed mouse manipulations including plasmid hydrodynamic injections, AAV injections, and serum and animal tissue collection. N.A. provided assistance with construct design and GUIDE-seq library preparation. A.M. analyzed deep-sequencing and GUIDE-seq datasets. R.I., A.M., and E.J.S. wrote the manuscript, and all authors edited the manuscript.

---

**Supplementary Figure 1:**
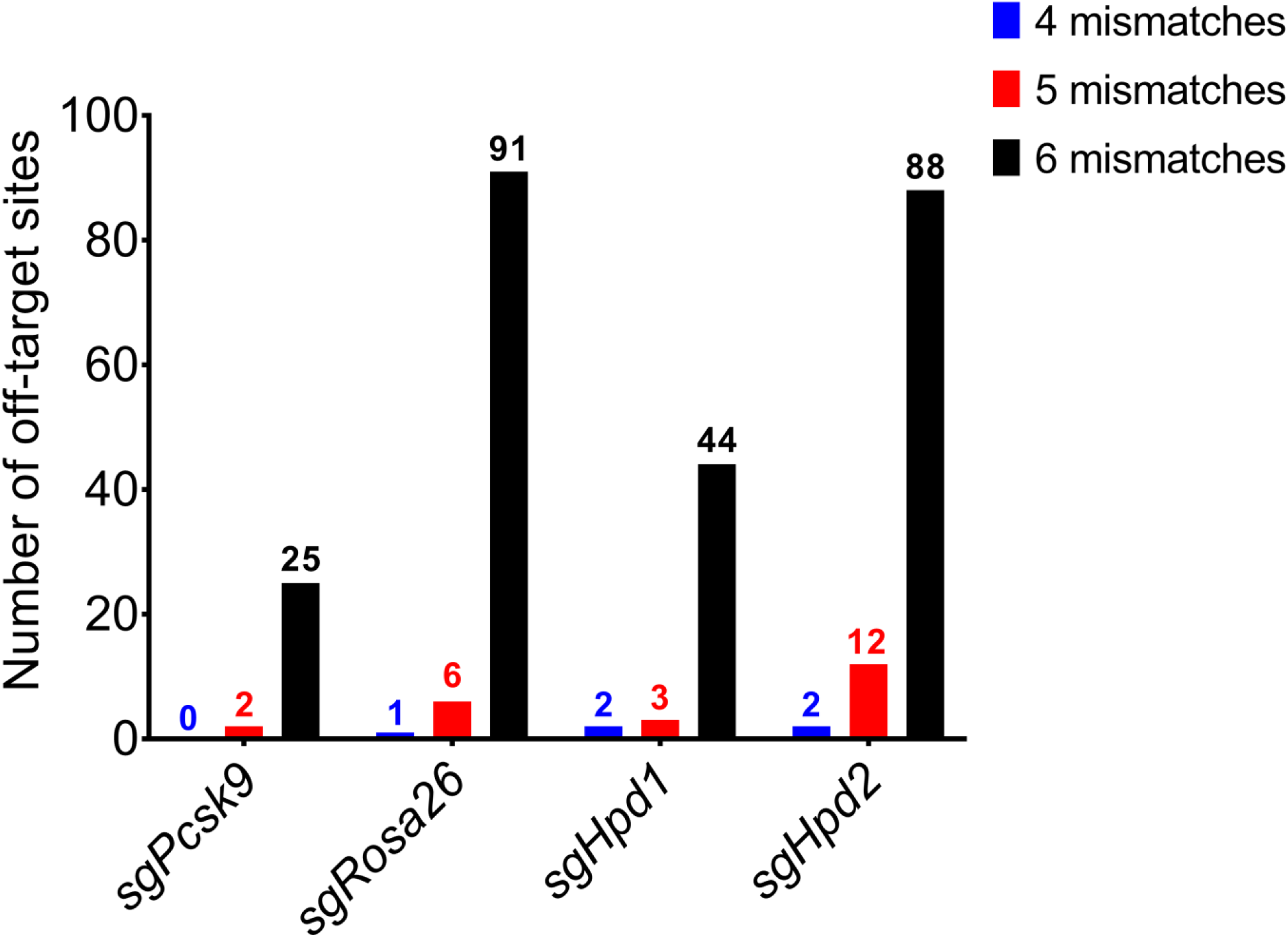
Genome-wide computational prediction of NmeCas9 off-target sites using CRISPRseek with the N_4_GN_3_ PAM. Search parameters were set to identify sites with up to 6 mismatches to the spacer sequence. The total number of detected off-target sites for each protospacer is indicated.

**Supplementary Figure 2:**
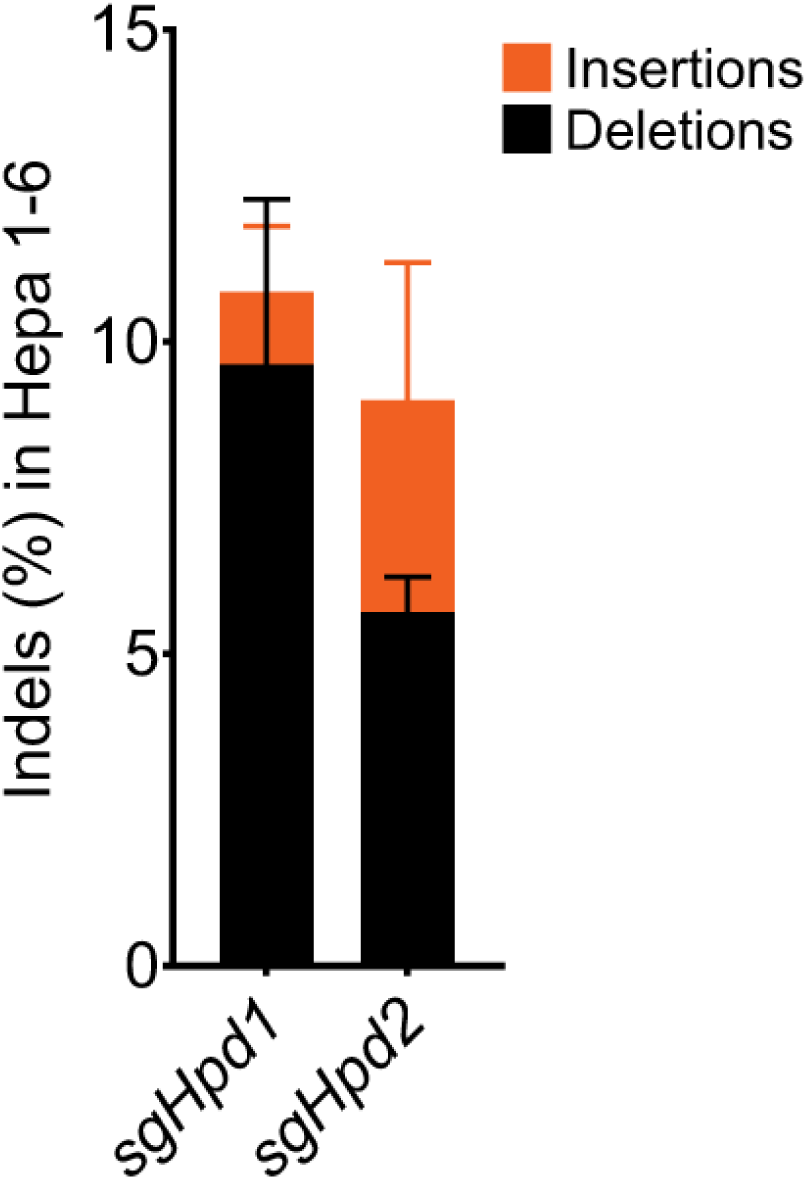
Stacked histogram showing the percentage distribution of indels obtained by TIDE after AAV-sgRNA-hNmeCas9 plasmids transfections in Hepa 1–6. Error bars represent three independent experiments. Data are presented as mean ± SD.

**Supplementary Figure 3:**
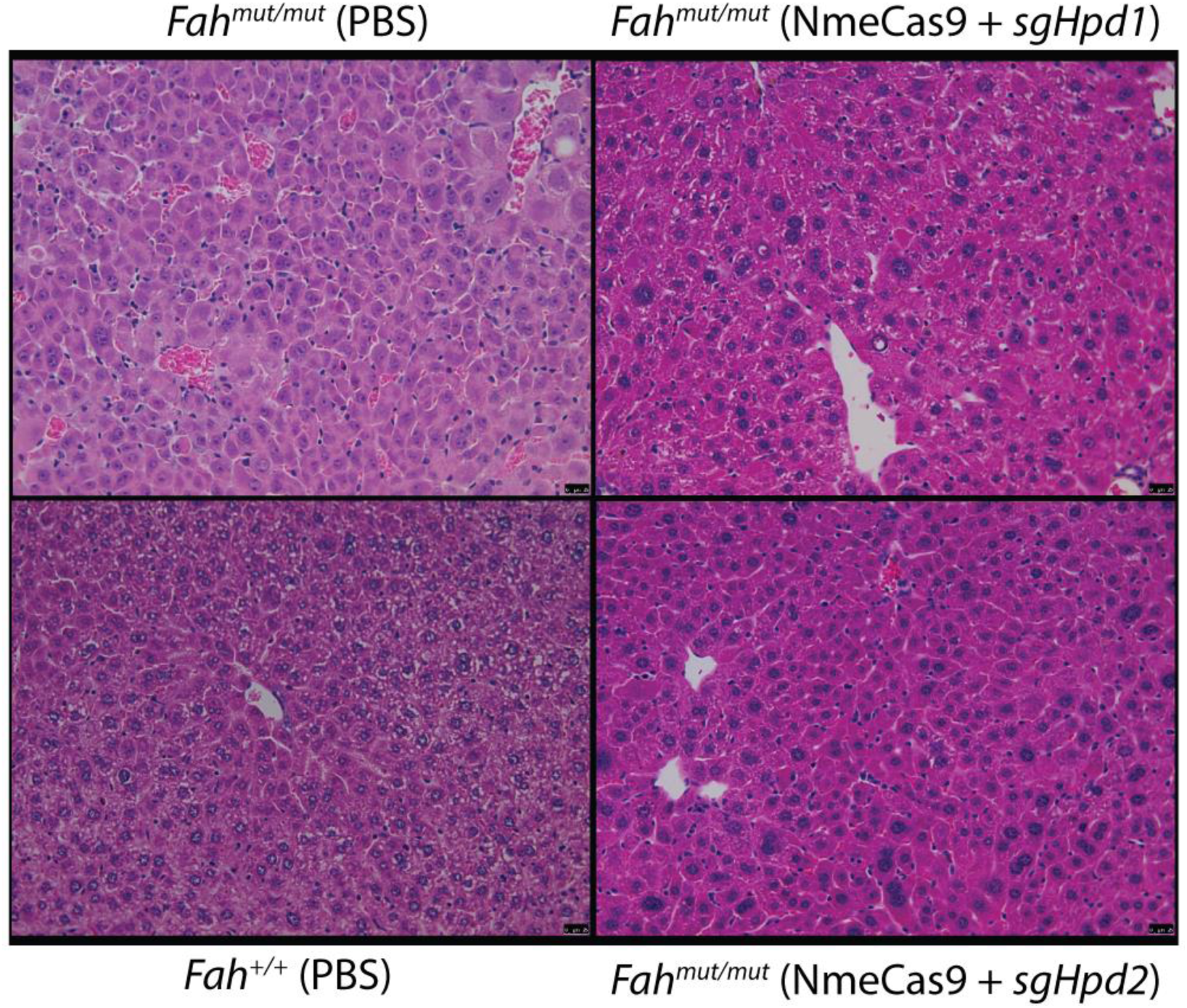
H&E staining from wild-type (*Fah^+/+^*) mouse, and HT-I mice (*Fah^-/-^*) injected with PBS or AAV-sgRNA-hNmeCas9 plasmids *sgHpd1* or *sgHpd2*. Scale bar is 20 μm.

**Supplementary Figure 4:**
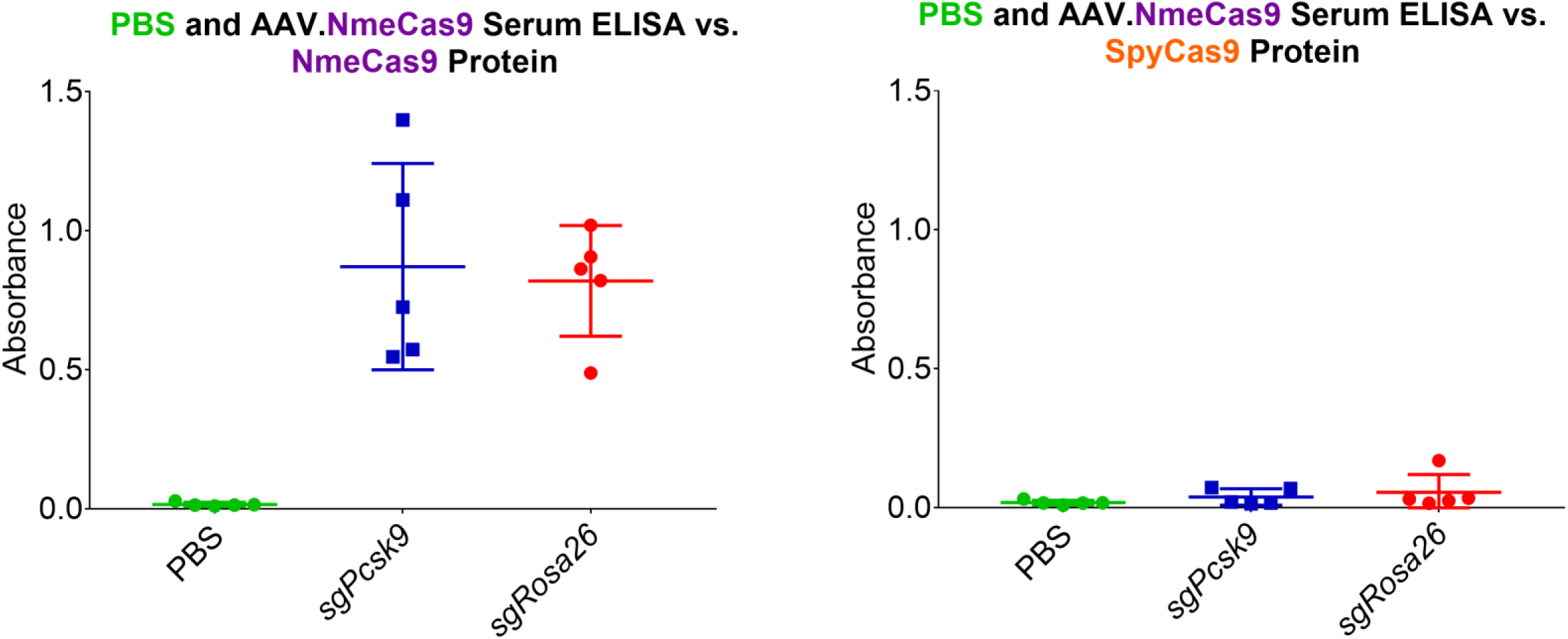
Humoral IgG1 immune response to NmeCas9 in vivo. Serum collected at day 50 post-injection with all-in-one AAV8-sgRNA-hNmeCas9 *sgPcsk9* and *sgRosa26*. Serum antibodies reacted against NmeCas9 protein (left) or SpyCas9 protein (right).

**Supplementary Figure 5:**
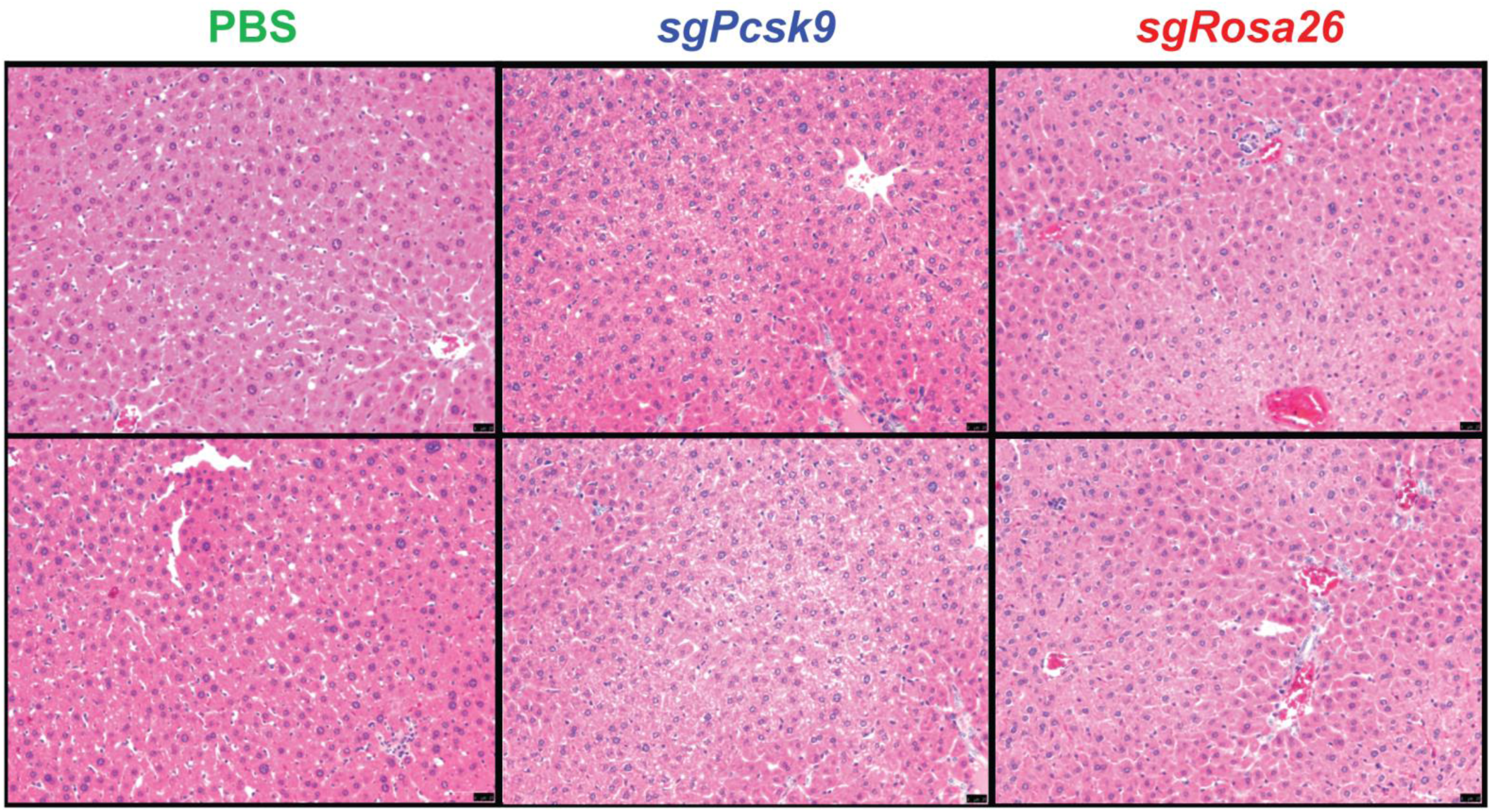
H&E staining from PBS, *sgPcsk9* and *sgRosa26* AAV8 injected mice. Scale bar is 20 μm.

## SUPPLEMENTARY TABLE

Protospacer sequences

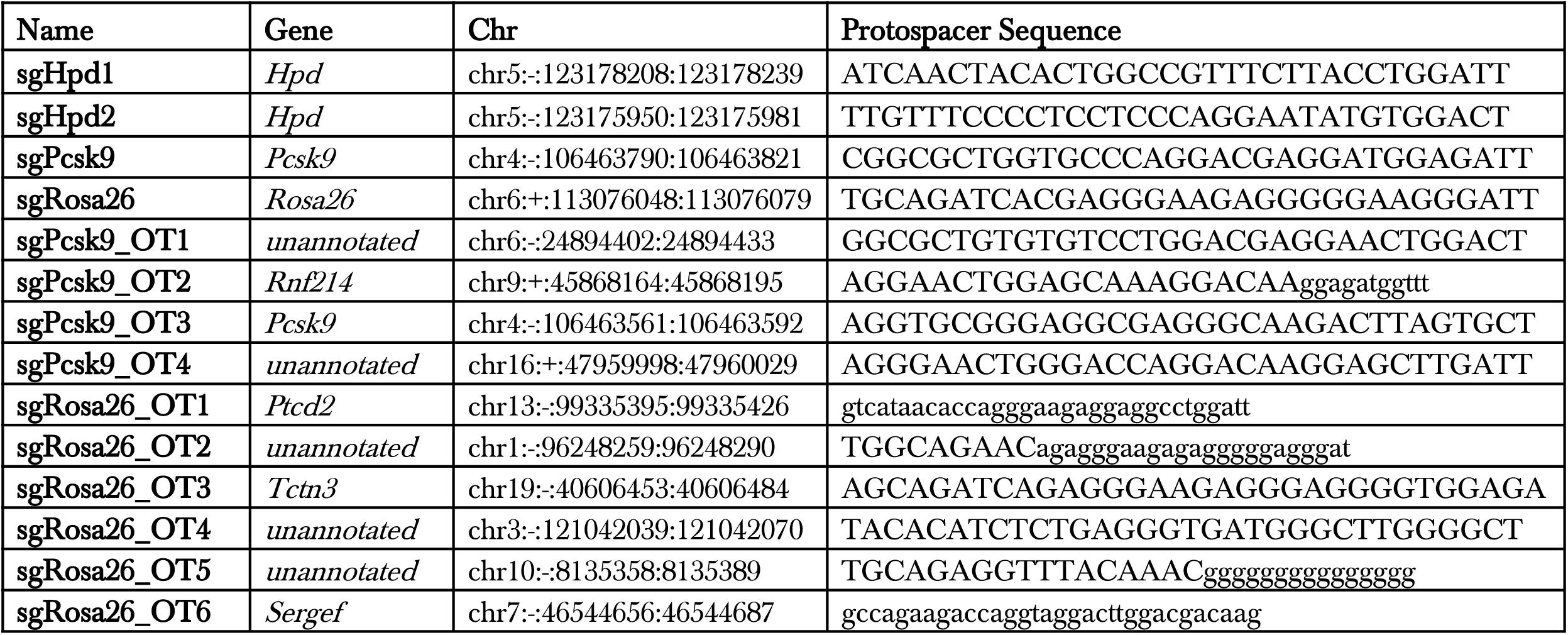

TIDE Primers

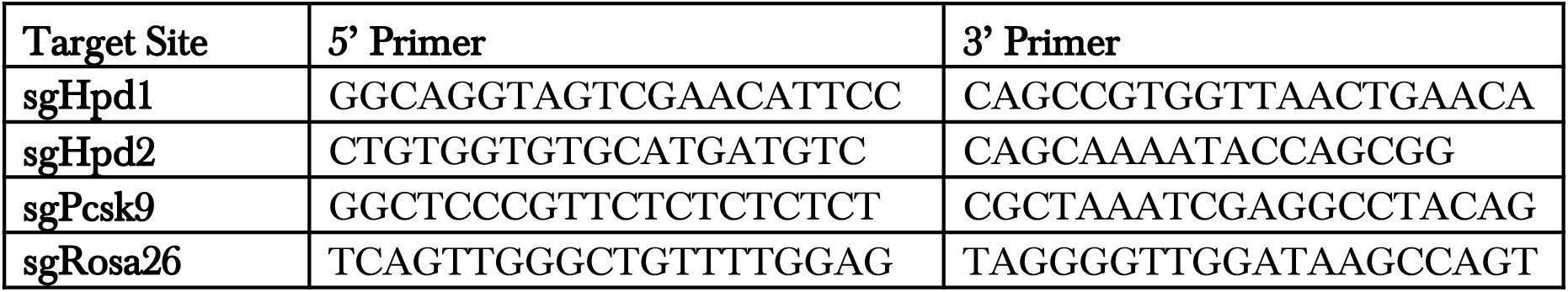

Deep-sequencing Primers

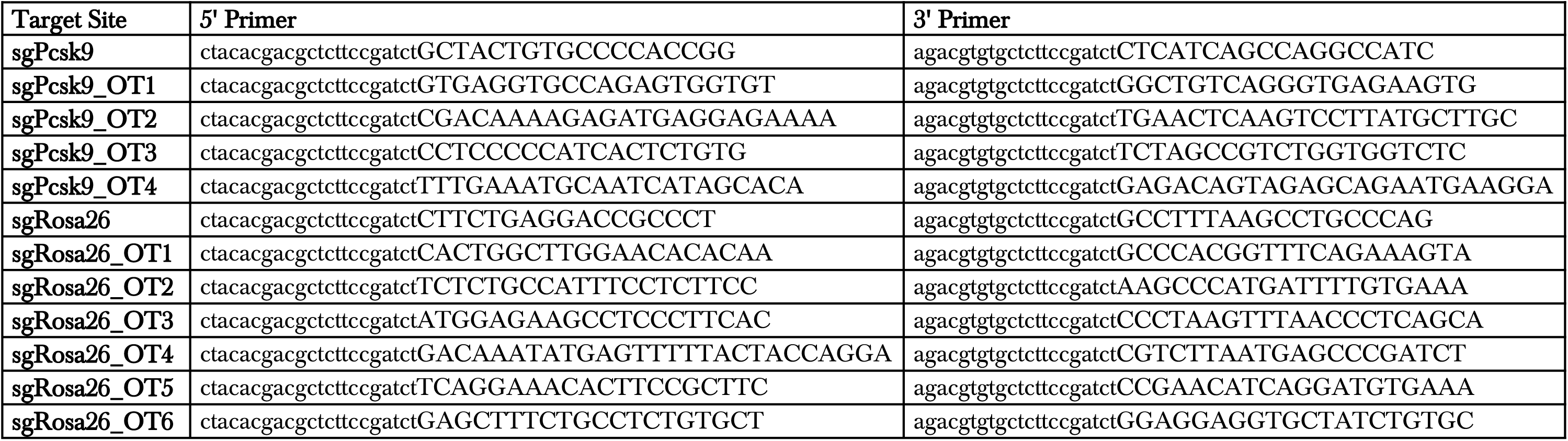

## SUPPLEMENTARY NOTE

Nucleotide sequence of all-in-one AAV-sgRNA-hNMeCas9 Plasmid

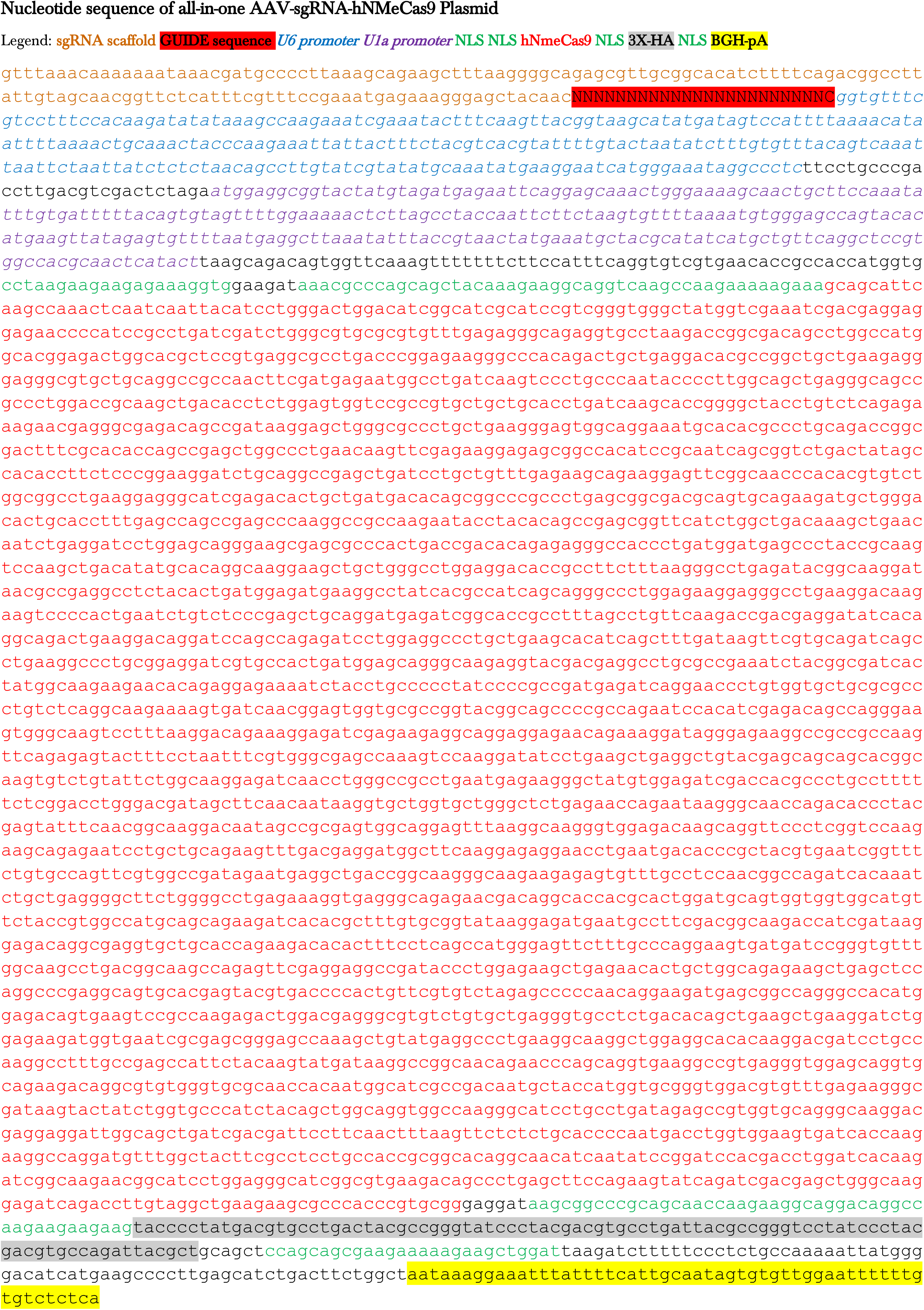

